# Experimental Kinetic Mechanism of P53 Condensation-Amyloid Aggregation

**DOI:** 10.1101/2025.02.03.635948

**Authors:** Silvia S. Borkosky, Ramon Peralta-Martinez, Alicia Armella-Sierra, Sebastián A. Esperante, Leonardo Lizárraga, Javier García-Pardo, Salvador Ventura, Ignacio E. Sanchez, Gonzalo de Prat-Gay

## Abstract

The tumor suppressor p53 modulates the transcription of a variety of genes constituting a protective barrier against anomalous cellular proliferation. High frequency “hot-spot” mutations result in loss-of-function by the formation of amyloid-like aggregates that correlate with cancerous progression. We show that full-length p53 undergoes spontaneous homotypic condensation at sub-micromolar concentrations and in the absence of crowders, to yield dynamic coacervates that are stoichiometrically dissolved by DNA. These coacervates fuse and evolve into hydrogel-like clusters with strong thioflavin-T binding capacity, which further evolve into fibrillar species with a clearcut branching growth pattern. The amyloid-like coacervates can be rescued by the HPV master regulator E2 protein to yield large regular droplets. Furthermore, we kinetically dissected an overall condensation mechanism which consists of a nucleation-growth process by sequential addition of p53 tetramers, leading to discretely-sized and monodisperse early condensates followed by coalescence into bead-like coacervates that slowly evolve to the fibrillar species. Our results suggest strong similarities to condensation-to-amyloid transitions observed in neurological aggregopathies. Mechanistic insights uncover novel key early and intermediate stages of condensation that can be targeted for p53 rescuing drug discovery.

**SIGNIFICANT STATEMENT:** Known as “the guardian of the genome”, the tumor suppressor protein p53 becomes activated by injuries to the DNA genome, and determines whether the cell must undergo self-destruction to avoid cancerous proliferation. P53 is in fact inactivated by mutations in over 50% of all cancers, and restoring its function is recognized as a therapeutic cancer target. A recent biochemical revolution in cell physiology and pathology are liquid entities known as biomolecular condensates. We show that p53 form condensates en route to pathological forms in a surprisingly similar manner to neurological amyloid diseases such as Alzheimeŕs and Parkinsońs. We uncover the sequence of steps in the reaction, exposing flanks for a novel drug development platform based on the condensates paradigm.

## INTRODUCTION

The tumor suppressor protein p53 constitutes a protective barrier against the initiation of cancer (1, 2). In normal cells, p53 levels are kept low by constitutive proteasomal degradation (2). In response to oncogenic or other types of cellular stress, p53 levels increase and modulate the transcription of a variety of genes involved in multiple activities, where the most salient are cell cycle regulation, DNA repair, senescence, and apoptosis (1, 2). The structural organization of p53 reflects the protein functional complexity, where active p53 is a tetramer with a modular domain structure consisting of independently folded DNA-binding and tetramerization domains, flanked by N- and C-terminal intrinsically disordered domains, adopting a highly dynamic conformational ensemble (3–7). P53 is able to modify this ensemble in response to environmental stimuli, adopting specific conformations compatible with interaction with a myriad of cellular partners (4, 8).

In over 50% of human cancers p53 is directly inactivated by mutation, with most mutational hot spots located within the DNA binding domain (DBD) (9–11). Depending on their location, these hot-spot mutations either decrease p53 thermodynamic stability or affect DNA-binding activity (9, 12). In the cellular context, these mutations can impact on the functionality of p53 either leading to a loss-of-function (LOF) phenotype, a dominant negative (DN) effect on the remaining wild-type p53, or even a gain-of-function (GOF) activity (12–14). Moreover, based on the central role of conformational instability of mutated p53 and consequent aggregation on cancer development, several small molecule and peptide inhibitors of aggregation designed to restore p53 native conformation have been explored as promising therapeutic strategies (15–18).

Studies that focused on the aggregation pathway of p53, showed co-aggregation between unstable hot-spot mutants and the wild-type protein in cell transfection assays (19). Further, *in vitro* characterization of this hetero-aggregation mechanism, proposed simultaneous unfolding, cross-reaction and cross-aggregation, rather than the classical seeding observed in prion-like processes (20). On the other hand, the irreversible aggregation of the independently folded p53DBD was shown to give rise to different morphologies, including amorphous, fibrillar, and prion-like, depending on physicochemical solvent conditions (21, 22). Despite growing evidence on cellular and tissue models that associate p53 amyloid formation with LOF and GOF phenotypes (23–25), the mechanistic aspects and characterization of these species for both wild-type and cancer p53 mutant is still scarce. Most of the *in vitro* studies with pure components have focused mainly on the p53DBD (21, 22), a monomeric species that covers only 50% of the protein sequence.

Biomolecular condensation by liquid–liquid phase separation (LLPS) underlies the formation of membrane-less organelles providing a mechanism for dynamic compartmentalization of the intracellular space (26–28). Typically, LLPS is promoted by hydrophobic and electrostatic interactions driven by intrinsically disordered regions (IDRs) of proteins, weak multivalent interactions, nucleic acid binding and, very importantly, the ability to oligomerize (29, 30). Moreover, condensates composed of proteins rich in IDRs are known to mature from liquid-like to solid-like states, particularly those related to neurological aggregopathies (26, 31). Recently, a multimutated p53 variant (17 amino acids, including all cysteines, except those involved in zinc coordination) was shown to undergo homotypic LLPS, proposed to be mediated by electrostatic interaction between the N-terminal transactivation domain (TAD) and the C-terminal regulatory domain (CTD), both intrinsically disordered (32). Under strong crowding conditions the p53DBD was shown to phase-separate *in vitro* and progressed into amyloid-like aggregates, and condensate-like GFP-p53 was reported in the nucleus (33). Another report described anomalous and amorphous wild-type p53 homotypic condensates, whereas the hot-spot mutant p53 R248Q formed mesoscopic protein-rich clusters, both hosting the nucleation of p53 amyloids (34, 35). To date, a handful of studies have explored phase separation on p53 (32–34, 36–42), and a phase separation-aggregation transition was proposed for p53 but not fully determined (33, 34). We have previously characterized the heterotypic LLPS process mediated by the interaction of p53 with the C-terminal domain of the E2 protein (E2C) of human papillomavirus 16 (HPV16), related to the HPV replication repression caused by p53 (36). However, the role of p53 in phase separation including a possible role in the amyloid aggregation pathway has not been established.

In this work we show that full-length pseudowild-type p53, but not monomeric p53DBD, spontaneously undergoes homotypic LLPS finely tuned by electrostatic interactions, where the regulatory C-terminal domain plays a main role. Although the N-terminal transactivation domain is dispensable for LLPS, it is a key participant in the on-pathway to amorphous aggregates and amyloid-like structures. While DNA reshapes and dissolves the condensates, the functionally linked HPV master gene regulator E2 can reversibly modulate the p53 aggregation route. In addition, we were able to i) experimentally address the fundamental condensation kinetic mechanism, ii) show that condensation is on-pathway to amyloid aggregation, and iii) show the formation of fibers with a clear branching pattern. We discuss the biological implications and possible impact on p53 rescuing therapeutics.

## MATERIALS AND METHODS

### Cloning

Specific primers were designed to generate p53ΔTAD, and p53ΔCTD truncated variants of p53 stable mutant (43) lacking the N-terminal transactivation domain (TAD) and the C-terminal domain (CTD), respectively. The PCR products were digested with BamHI and EcoRI enzymes, purified and inserted into a modified pRSETa vector in frame with an N terminal fusion 6xHis/lipoamyl domain/TEV protease cleavage sequence. Sequencing the entire coding regions and flanking sequences confirmed the truncations of the full-length protein.

### Expression and purification of recombinant proteins

The pET24a plasmids containing the sequence of stable mutants of full-length p53 (FL-p53), and DNA-binding domain (p53DBD) with the following mutation: M133L/V203A/N239Y/N268D(43) was a kind gift from Alan Fersht. These mutations conserve the functional and conformational properties of wild-type p53, allowing increased levels of expression and sample stability. Therefore, it is considered as pseudo wild-type p53 protein. Pure FL-p53 and p53ΔTAD were obtained as previously described (36). P53ΔCTD and p53DBD were expressed in *Escherichia coli* C41 strain and purified by using a standard His-tag purification protocols, followed by tobacco etch virus (TEV) protease digestion. In the case of p53ΔCTD an ionic exchange QHyperD column was used, whereas for p53DBD a second His-tag purification step, allowed further purification. The final purification step was size exclusion chromatography (SEC) using Superdex-200 column (Cytiva) for tetrameric p53ΔCTD, and Superdex-75 column (Cytiva) for monomeric p53DBD. p53 conformations was confirmed by estimating the hydrodynamic radius by dynamic light scattering (DLS). The HPV-16 E2 C-terminal domain (E2C) was expressed in *Escherichia coli* BL21(DE3) and purified as described previously (44, 45). Protein concentration was determined by UV light absorbance at 280 nm using the protein molar extinction coefficients and confirmed by Bradford assays. Purified proteins were stored as at −80 °C after snap freezing in liquid nitrogen and aliquoted in fixed volumes to avoid repetitive thaw-freezing. Far-UV circular dichroism and fluorescence spectra, and SDS-PAGE were performed to confirm the quality and purity of the proteins, respectively.

### Fluorescent labeling

FL-p53 and E2C proteins were labeled with fluorescein isothiocyanate (FITC) (Sigma-Aldrich), adapting the manufacturer’s protocol to obtain sub-stochiometric labeling enough to visualize the samples by fluorescence and confocal microscopy. A p53/FITC ratio of 2 and a E2C/FITC ratio of 6 were used. Reactions were carried out at 4 °C overnight in potassium phosphate buffer 50 mM, NaCl 0.3 M and 1 mM DTT pH 7.0. The reactions were stopped using 50 mM Tris-HCl pH 8.0 and excess of FITC removed by desalting PD10 columns (G&E) eluting each protein with the corresponding stock buffers. A similar procedure was carried out for labeling all p53 variants with cy5 NHS (Lumiprobe) using a p53/cy5 ratio of 2. These protocols yield proteins stocks labelled with 10% -20% fluorescent dye.

### DNAs

Double-stranded 26 bp oligonucleotides containing p53 consensus sequence (46) were prepared as follows: single-stranded oligonucleotides were purchased, HPLC purified, from Integrated DNA Technologies (Coralville, IA). p53_DNA_A: 5‘ AGC TT *AGGCATGTCT AGGCATGTCT* A 3° and p53_DNA_B: 5‘ AGC TT *AGACATGCCT AGACATGCCT* A 3° (Recognition sequences are italicized). Single-stranded oligonucleotide concentration was calculated using the molar extinction coefficient obtained from the nucleotide composition. Annealing was performed as described previously(44). This yielded double-stranded oligonucleotide termed DNA_p53_, and no detectable single-stranded oligonucleotide was judged by PAGE. Calf thymus DNA (ctDNA) (Sigma-Aldrich) was dissolved following the manufacturer’s instructions to obtain a 1 mg/ml stock solution. ctDNA quantification was confirmed by nanodrop (Thermo Scientific) and quality was assessed by agarose gel electrophoresis.

### Bright field microscopy and fluorescence microscopy

Analyses of homotypic p53 LLPS and heterotypic p53/E2C LLPS were performed using bright field and florescence microscopy, under varying conditions, including concentration, ionic strength, and molecular crowding. Samples were prepared at room temperature using 0.3 μM labeled protein within the bulk of unlabeled materials and loaded into 96-well plates (Corning nonbinding surface). Sample buffer consisted of 50 mM Tris-HCl buffer, 40 mM and 1 mM, DTT pH 7.0. When molecular crowding conditions were needed, sample buffer consisted in 50mM Tris-HCl buffer, 150 mM NaCl, 1 mM DTT and 10% PEG, pH 7.0. For fluorescence analysis, the samples were excited at 488 nm for FITC or 633 nm for cy5. The images were acquired using an Axio Observer 3 inverted microscope with a 40x/0,750.3 M27 objective and a Colibri 5 LED illumination system. Images were processed using Fiji (A distribution package of ImageJ software, USA) (47). For thioflavin T (ThT) staining, 50 µM of the ThT (Sigma-Aldrich) was included in the sample buffer. All reagents were filters through a 0.2 µm filter prior to use.

### Fluorescence recovery after photobleaching

2.5 µM FL-p53 samples were prepared and imaged using Nunc Lab-Tek chambered coverglass (ThermoFisher Scientific Inc) pre-coated with 1% Pluronic F-127 (Sigma-Aldrich) at room temperature, with 0.3 μM labeled samples within the total of protein concentration. Fluorescence recovery after photobleaching (FRAP) experiments were performed using a Zeiss LSM 880 Airyscan confocal laser scanning microscope with a Plan-Apochromat 63/1.4 objective lens as described previously (36). Fluorescence recovery data were evaluated using the FIJI ImageJ software(36). Data was fit to a two-exponential fit using equation 1 *y* = *A1*\*exp(−*k_1_*t*) + *A2*exp*(−*k_2_*t*), where *A* and *k* are fitting parameters. Half time of recovery (*t*_1/2_) was obtained graphically.

### Turbidity measurements

Protein samples were prepared in 50 mM Tris-HCl buffer, 20-170 mM NaCl (depending on the salt concentration needed), and 1 mM DTT, pH 7.0 or 50 mM Tris-HCl buffer, 150 mM NaCl, 1 mM DTT, and 10% PEG, pH 7.0, in a total volume of 100 μl. The p53 concentration ranged from 0.25 to 10 µM. For FL-p53, p53ΔTAD, and p53ΔCTD concentration is expressed in tetramer, whereas for p53DBD is referred as monomer. The samples were incubated at room temperature for 15 min, and absorbance at 370 nm was measured using a Varioskan LUX 3020 (ThermoFischer Scientific). All the conditions were measured in 96-well plates (Corning nonbinding surface). Each turbidity assay was reproduced in triplicate. All reagents were filters through a 0.2 µm filter prior to use.

### Thioflavin T binding assays

Fluorescence emission spectra were recorded on a Horiba FluoroMax 4 spectrofluorimeter with an excitation wavelength of 450 nm, at protein tetramer concentration of 2.5 µM in 50 mM Tris-HCl buffer, 50 or 150 mM NaCl, 1mM DTT, 25 µM ThT, pH 7.0 at 25 °C. All reagents were filters through a 0.2 µm filter prior to use.

### Turbidity kinetic measurements

p53 condensation was recorded in a Jasco UV spectrophotometer by following scattering signals at 370 nm, using 700 µl volume in a 1 cm optical path quartz cuvette. Measurements were performed in 50 mM Tris-HCl 40 mM - 200 mM NaCl (depending on the condition) and 1 mM DTT, pH 7.0. When crowding was needed, 5% PEG-4000 was used. Measurements were conducted in different samples containing p53 concentrations that ranged between 0.25 μM and 2.5 μM), referring always to tetramer concentration.

For mechanistic analysis, turbidity kinetic traces were fitted using a two-exponential function plus a drift, using the following equation 2 *y* = *A1*\*exp(*k_1_*t*) – *A* exp*(*k_2_*t*))+m *t*, where A and *k* are fitting parameters. Next, NAGPKin Web Server was used applying the model described by Martins and coworkers (48).

For the nucleus size determination we used the KL model (49), in which the relationship between the concentration of the condensed p53 and the free protein at a certain time during the assembly is given by the equation 3: [p53cond] = *k*\*[p53free]*^n^*. This relationship is valid early in the reaction and before reaching the steady state. In equation 3, *k* is a proportionality constant, and *n* reports the nucleus size. The nucleus size *n* is calculated from the slope of a log-log plot of condensed p53 and p53 free molar concentration. Each straight line was obtained calculating the p53 condensed and p53 free concentrations obtained at a single time for different initial total protein concentrations. P53 free concentration is tetrameric concentration. This model was applied in the initial turbidity rates by fitting a straight line to the first 110 seconds of the process. The turbidity signal is proportional to the condensed p53 assembly: [p53cond]∼A_370nm_ signal. The free protein was taken from equation 4: [p53free]∼[p53]initial*(1-A_370nm_ signal).

### Dynamic light scattering

DLS measurements were carried out on DynaPro NanoStar II DLS device (Wyatt Technology). Measurements were performed in 50 mM Tris-HCl, 40 mM or 150 mM NaCl, DTT 1 mM, pH 7.0. p53 samples were prepared at a fixed tetramer concentration of 1.25 µM. The temperature was maintained at 25°C by a Peltier control system. Results were processed employing the software package included in the equipment. The kinetics assay was carried out by averaging a set with 10 measurements with a read interval of 1 sec, an acquisition time of 20 seconds, for 4500 seconds. DLS trace was fitted using a single-exponential function using the equation 5 *y* = *A1*\*exp(*k_1_*t*)+m\**t*.

### Transmission electron microscopy

For negative Staining Transmission Electron Microscopy (NS-TEM), 10 µl of each sample were placed on top of a EMR 400 mesh carbon-coated copper grids (Micro to Nano Innovative Microscopy Supplies) and incubated at room temperature for 5 min. After incubation, the grids were stained for 1 min with 5 µl of 2% (w/v) uranyl acetate. The excess of sample and staining solutions were removed using filter paper. After staining, the grids were allowed to dry before inspection using a JEOL JEM-1400 Electron Microscope at 120 kV. The microscope was equipped with a CCD GATAN 794 MSC 600HP camera controlled by Digital Micrograph 1.8 (GATAN) software. 3–5 representative micrographs were recorded for each sample at the indicated incubation times. The estimated defocus of the images was about ± 1– 5 μm. Image analysis and size measurements were carried out with ImageJ software (47).

### Atomic force microscopy

For the sample preparation, a volume of 40 µL of each sample were deposited on a muscovite mica discs grade V1 (Ted Pella, Inc.) of 10 mm diameter, which were previously cleaved using a scotch tape and glued to steel discs. Then, the discs were dried for 24 hours at room conditions to achieve total dryness. Experiments were carried out in a temperature-controlled room at 25 ± 1 °C, with acoustic hood isolation and active vibration damping. The AFM images were acquired in tapping mode using silicon tips with a spring constant of 42 N m^−1^ and a resonance frequency of 320 kHz, using a Bruker Multimode 8 SPM (Santa Barbara, CMA, USA) and a NanoScope V Controller (Santa Barbara, CMA, USA). The image analyses were performed using Gwyddion version 2.46 software (Brno, Czech Republic).

## RESULTS

### Condensation of p53 is finely modulated by ionic strength and strongly dependent on its C-terminal regulatory domain

Human p53 is a highly stable tetramer (50, 51) with a complex domain organization, consisting of four domains: i) acidic intrinsically disordered N-terminal transactivation (TAD) including a proline rich region, ii) globularly folded DNA-binding (DBD), iii) a folded tetramerization (TET), and iv) C-terminal DNA binding modulatory (CTD) (Figure 1A). Although the DBD has mostly positively charged amino acids, it also contains a considerable number of negative residues (Figure 1A). Wild-type full-length p53 is known for a marked tendency to readily undergo thermal aggregation under mild physiological conditions (43), which hampers purification and biochemical manipulation. In order to overcome this, we used a rationally designed stability mutant of the full-length protein, previously described as pseudowild-type p53 (FL-p53), which is indistinguishable from wild-type p53 in terms of tetramer assembly and DNA binding function (43).

**Figure 1.**
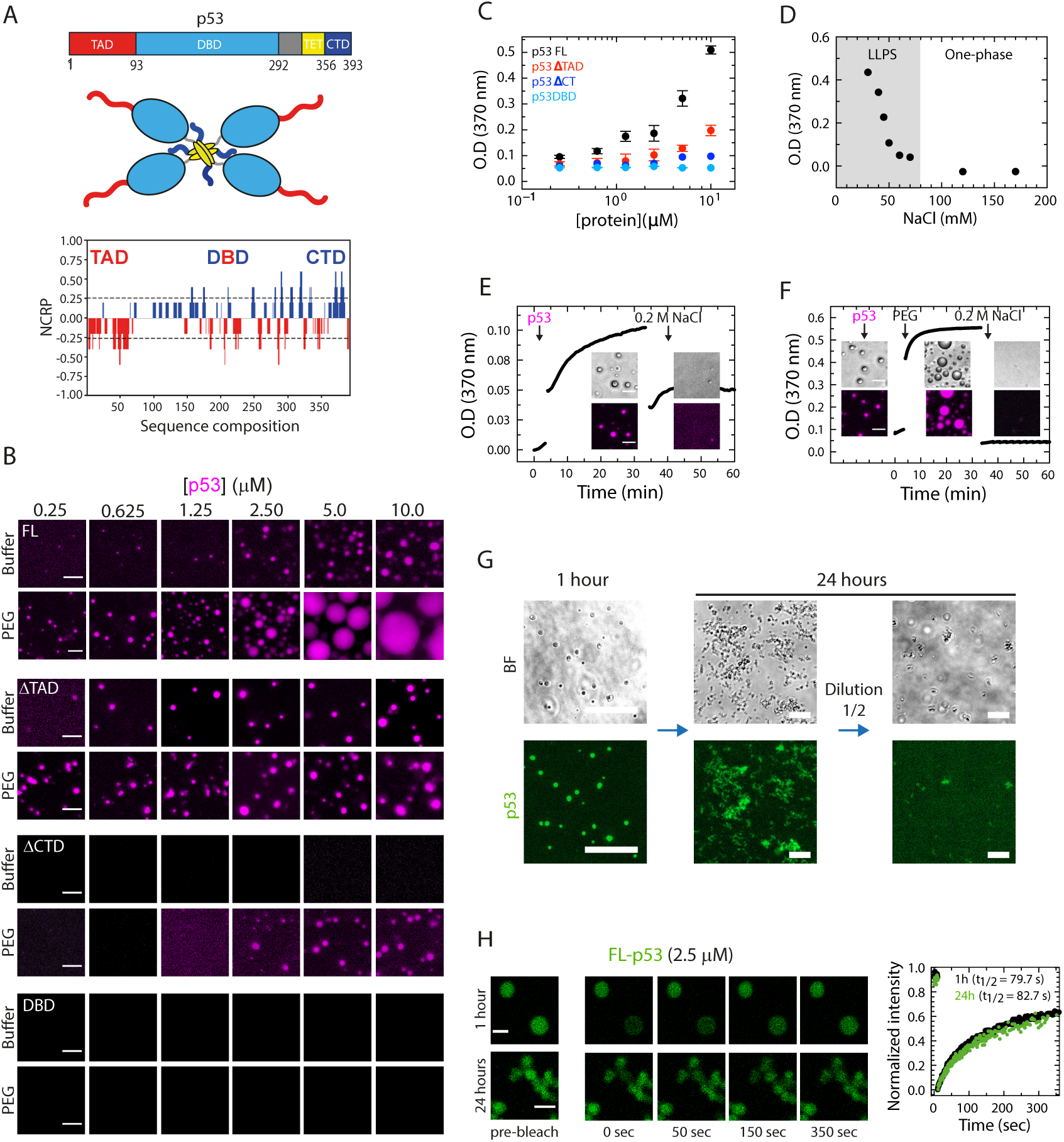
p53 forms spherical droplets that depend on its C-terminal domain and are modulated by ionic strength. (A) Top, Schematic organization domain of full-length wild-type p53. Middle, schematic representation of the p53 structure showing the tetrameric conformation of the protein with each domain highlighted in different colors. Bottom, charge patterning of the p53 full-length sequence using CIDER (http://pappulab.wustl.edu/CIDER/analysis/) (52). NPCR, net charge per residue. (B) Representative images of samples with increasing concentrations of cy5-labaled full-length p53 (FL-p53) and p53 truncated mutants, lacking the N-terminal TAD domain (ΔTAD), the C-terminal regulatory domain (ΔCTD), and the monomeric p53 DNA-binding domain (DBD). Bar on images= 10 µm. (C) Turbidity assay monitoring absorbance at 370 nm on individual samples containing 2.5 μM FL-p53 (Black circles), p53ΔTAD (red circles), p53ΔCT (blue circles) and p53DBD (turquoise circles) in presence of 10% PEG-4000. (D) Turbidity assay monitoring absorbance at 370 nm of individual samples with increasing concentrations of NaCl. Grey background indicates the salt concentration range permissive for LLPS, white background indicates the system is under a one-phase regime. (E) Kinetic turbidity assay of a sample containing cy5-p53 monitored by light scattering combined with visualization by bright field and fluorescence microscopy; NaCl was added after the signal reached maximum absorbance. Bar on images= 10 µm. (F) Similar experiment as in E, but in presence of crowding agent (5% PEG-4000). Bar on images= 10 µm. (G) Representative images of 2.5 µM FITC-FL-p53 incubated for 1 hour (left panel), 24 hours (middle panel) and samples incubated for 24 hours, and then diluted (right panel). BF, bright field. Bar on images= 20 µm. (H) Representative confocal microscopy images and mean plot of fluorescence recovery after photobleaching (FRAP) analyses of homotypic condensates composed of 2.5 μM FITC-labeled p53 (FITC-p53) incubated for 1 hour (n= 7 droplets) or p53 coacervates incubated for 24 hours (n= 5). Data were normalized to the average intensity of a droplet/region not photobleached and fitted using a double exponential function. Half time recovery (*t_1/2_*) of 79.7 ± 11.5 or 82.7 ± 34.6 seconds were obtained for 1 hour and 24 hours incubated samples, respectively. Scale bars= 2 μm (upper panel) and 3 µm (lower panel).

Upon dilution of FL-p53 into a low ionic strength buffer we observed small regular droplets by both fluorescence and bright field (BF) microscopy (Figure 1B and Figure S1A). The size of the droplets increased with the increase in protein concentration over the range tested, reaching a maximum diameter of 3 µm in the absence and 30 µm in the presence of PEG as crowding agent (Figure 1B and Figure S1A). The onset for the formation of the condensates under crowding conditions is ∼0.6 µM compared to ∼2.5 µM FL-p53 tetramer concentration in the absence of PEG (Figure 1B and Figure S1A). Control experiments confirmed that droplet formation was independent of the type or ratio of fluorescent label used (Figure S1B). In order to determine which regions of p53 were involved, similar *in vitro* LLPS assays were carried on by using p53ΔTAD and p53ΔCTD deletion mutants (Figure 1B and Figure S1A). The p53ΔTAD was able to form spherical droplets indistinguishable from those of FL-p53 (Figure 1B and Figure S1A) but the effect of the crowder was less accentuated than that observed for FL-p53 (Figure 1B and Figure S1A). Turbidity assays showed a lower tendency for LLPS by p53ΔTAD compared to FL-p53 (Fig. 1C). No condensate droplets were observed for p53ΔCTD in the absence of crowder up to 10 µM p53 tetramer (1.6 mg/mL) but spherical discrete condensates were found when PEG was added, in protein concentrations of 2.5 µM and above (Figure 1B and Figure S1A), suggesting a lower but significant tendency to LLPS. Finally, no evidence of condensate or aggregates was observed for p53DBD under similar condition to the FL and deletion variants of p53, either in absence of presence of PEG (Figure 1B and Figure S1A).

Given the ampholytic nature of p53 as evidenced in a net charge per residue (NCPR) plot (52) (Figure 1A) and previously observed ionic strength effect on multimutated variants (32), we wanted to evaluate how ionic strength modulates p53 condensation. Changes in turbidity signal at 370 nm as a function of NaCl concentration showed a sharp transition below 50 mM salt (Figure 1D). Time course experiments after transferring concentrated FL-p53 from ice to room temperature and low salt (40 mM NaCl) led to an increase in turbidity reaching a plateau at 30 minutes, which corresponded to the formation of the small regular droplets (Figure 1E). Subsequent addition of 200 mM NaCl completely reverted both the scattering signal and the formation of droplets (Figure 1E). A comparable experiment was performed in the presence of 5% PEG, with a more pronounced change in the absorbance scattering signal, corresponding to the formation of larger and highly regular spherical droplets, of up to 7 µm in diameter (Figure 1F). Addition of 200 mM NaCl also fully dissolved the droplets, indicating that droplet condensation is reversibly governed by ionic strength both in the presence and absence of a crowding agent.

We next aimed at addressing the evolution and properties of the FL-p53 condensates. To this end, these droplets displayed partial coalescence into non-regular clusters (Figure S1C), suggesting some degree of maturation over time. Indeed, the droplets evolved to clustered bead-like structures highly suggestive of hydrogel coacervates (Figure 1G). Following dilution, the 24-hour aged species were disassembled almost completely (Figure 1G). Moreover, fluorescence recovery after photobleaching (FRAP) experiments in 1 hour or 24-hour aged FITC-FL-p53 yield a ca. 70 % recovery, with a *t*_1/2_ of 80 seconds (Figure 1H and Figure S1D). Overall, the results indicate that the impossibility of undergoing complete coalescence is not related to internal properties of the condensates, most likely surface tension phenomena that lead to sticky bead-like clustered hydrogels.

### p53 condensates are on-pathway to amyloid-like fibrillar aggregates

Recent studies proposed that p53 condensates might nucleate the formation of amyloid-like structures (33, 34), but experimental evidence for on-pathway presence of the condensates is still lacking. In order to determine a sequential link between FL-p53 condensates and amyloid-like aggregates, we used thioflavin T (ThT) as a reporter of amyloid-like repetitive β-sheet structure formation. Time course experiments showed a clear fluorescence increase within the coacervates starting at 5-hour incubation, indicative of binding of ThT to p53 species different from the conformation in the dilute phase. These bead-like coacervates evolved into denser coacervates with strong ThT staining after 24 hours incubation (Figure 2A). After one week incubation, there was more pronounced ThT binding in the p53 aggregates, that occupied a wider area at the bottom of the plate (Figure 2A). In the presence of PEG, the droplets retained a highly regular spherical shape, increasing in size with time due to coalescence (Figure S2A). However, only a faint ThT staining was observed after 24-hour incubation in crowding conditions, with a more intense ThT staining after 1 week incubation (Figure S2A). This result suggests that in the presence of crowding agent a different p53 conformation is present (Figure S2A). Control experiments, without ThT, were performed in the absence or presence of PEG using either cy5-FL-p53 or FITC-FL-p53, displaying identical morphologies as when ThT was added (Figure S2B, and C). This result also showed that ThT binding does not interfere with the condensates formation and evolution. Next, ThT binding experiments in solution were carried out, showing a slight increase in the ThT signal between one hour and 24-hour incubation period (Figure 2B). Similar results were obtained when assessing ThT spectra of samples incubated for 1 week in comparison to samples incubated for 1 hour (Figure S2D). Control samples containing 150 mM NaCl (LLPS abrogation, Figure 1E and 1F) in the incubation buffer showed no difference in their ThT fluorescence spectra, in comparison with the LLPS samples in buffer with 40 mM NaCl (Figure 2B and Figure S2D). This result indicates that although a strong ThT binding signal could be detected in droplet coacervates (Figure 2A), only a marginal ThT fluorescence is observed in solution. A likely explanation is that ThT fluorescence will only be detected in soluble species, but not in early insoluble amyloid-like species.

**Figure 2.**
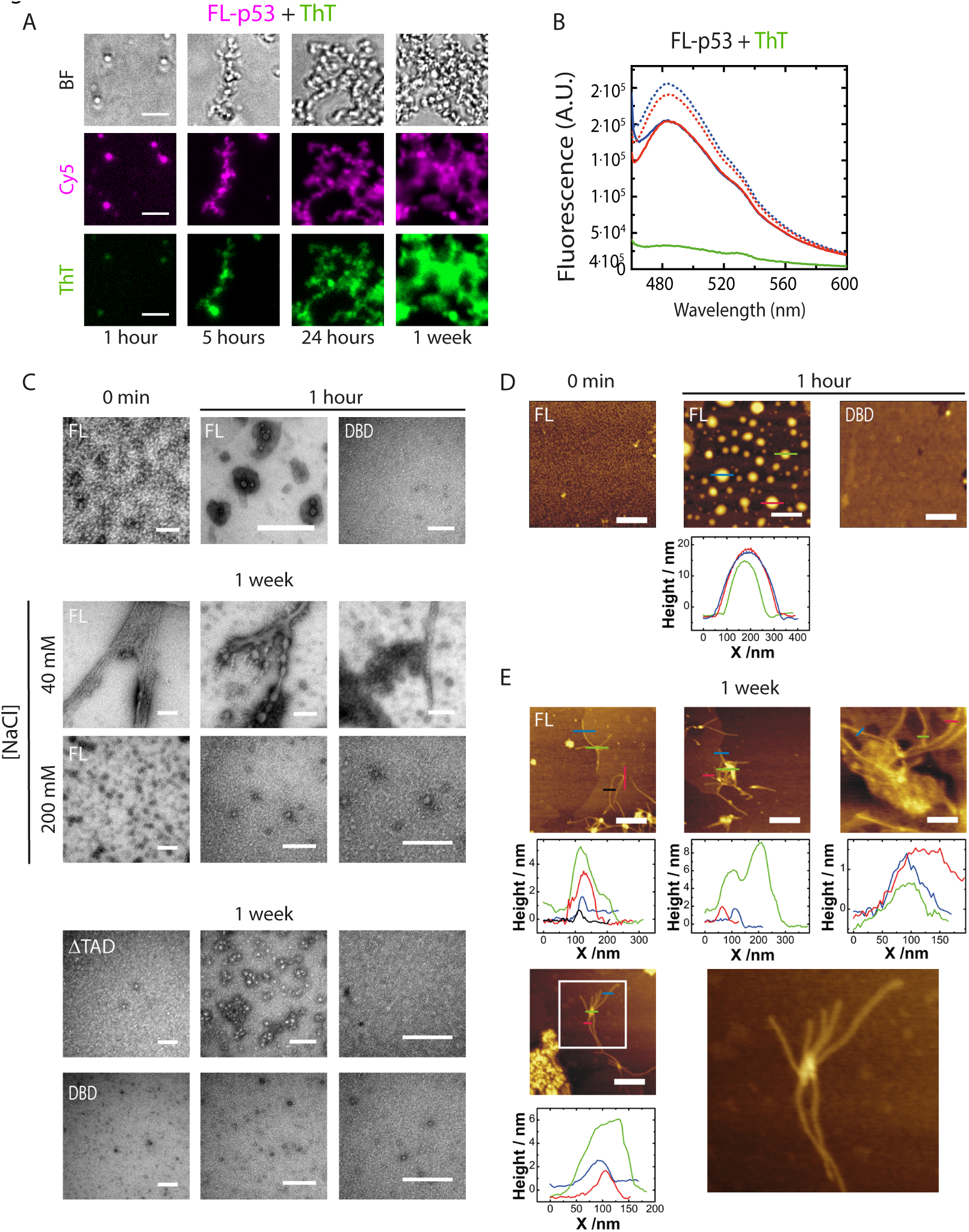
p53 droplets evolve to amyloid-like and amorphous aggregates. (A) Representative microscopy images of samples containing cy5-FL-p53 incubated with Thioflavin T (ThT) during 1, 5, 24 hours, and 1 week without crowding agent. BF, bright field. Scale bars= 10 μm. (B) Fluorescence spectra of FL-p53 in buffer containing 25 µM ThT and 40 mM NaCl (Blue) or 150 mM NaCl (Red), pH 7.0, incubated for 1 hour (Thick line) or 24 hours (Dotted line). The spectrum depicting the fluorescence emission of buffer containing ThT without protein is shown in green. (C) Top, representative transmission electron microscopy images (TEM) of FL-p53 visualized immediately after sample preparation (0 min) and images of FL-p53 and p53DBD after incubation for 1 hour in buffer containing 40 mM NaCl, pH 7.0. Middle, TEM images of FL-p53 samples incubated for 1 week in presence of 40mM or 200 mM NaCl. Bottom, TEM images of control samples containing p53ΔTAD and p53DBD incubated for 1 week in presence of 40 mM NaCl. Scale bars= 200 nm. (D) Atomic force microscopy (AFM) images of samples of FL-p53 visualized immediately after preparation (0 min) and images of FL-p53 and p53DBD incubated for 1 hour. Height profiles of FL-p53 droplets incubated for 1 hour are shown below the image. Scale bars= 500 nm. (E) AFM images with height profiles of FL-p53 samples incubated for 1 week. Frame on image is shown magnified for detailed visualization. Scale bars = 500 nm.

To evaluate the evolution of the amyloid-like material product of condensate maturation, we carried out transmission electron microscopy (TEM). Negative staining samples of FL-p53 incubated under LLPS conditions and visualized immediately after sample preparation showed the presence of soluble and oligomeric FL-p53 species (Figure 2C, top panel and Figure S3A). In contrast, after 15 min and 1 hour-incubation we observed droplet-like structures (Figure 2C, top panel and Figure S3A), which are compatible with the FL-p53 condensates previously observed by bright field and fluorescence microscopy (Figure 1B and Figure S1A). Noticeably, FL-p53 incubated for one week under LLPS conditions (low salt, i.e. 40 mM NaCl) showed fiber-like structures together with some amorphous aggregates (Figure 2C, middle panel and Figure S3A). In the presence of salt concentrations that abrogate p53 LLPS (*i.e.* NaCl 200 mM), no fiber-like material was observed after one week incubation (Figure 2C, middle panel, and Figure S3B), linking condensation with amyloid aggregation. The p53ΔTAD variant showed spherical condensates observed by low magnification microscopy (Figure 1B). When incubated for one week under LLPS and non-LLPS conditions, no fibrous material was observed by TEM (Figure 2C, bottom panel and Figure S3B). When p53DBD was incubated under the same experimental conditions we observed neither condensates at one hour incubation (Figure 2C, top panel) nor fiber-like material after one week incubation (Figure 2C, bottom panel and Figure S3B).

In order to complement our findings, we tackled an atomic force microscopy (AFM) analysis. Spherical droplets of approximately 200 nm diameter and a height of 15 nm could be observed within the first hour of incubation under LLPS (Figure 2D, and Figure S3C), in agreement with bright field and fluorescence microscopy (Figure 1B and Figure S1A). However, following one week incubation, mostly fibrillar morphologies of variable sizes were observed, with some amorphous aggregates (Figure 2E, and Figure S3C). Interestingly, a fibrillar branching pattern was observed, where branches project and grow from large nuclei-like structures with sizes ranging from 5 to 8 nm (Figure 2E). Samples of p53DBD incubated in parallel under identical conditions as FL-p53 showed formation of neither droplets nor aggregates (Figure 2D and Figure S3D).

### Kinetic mechanism of p53 condensation

To understand and dissect the homotypic p53 condensation mechanism from the native tetramer to the droplets, we tackled a detailed kinetic analysis. To this end, we followed the time course of changes in turbidity by monitoring absorbance at 370 nm, under the LLPS low salt favoring conditions, and at different initial p53 tetramer concentrations (Figure 3A). A slow change can be observed with no evidence of fast burst-phase events, indicating no condensation within the experimental deadtime (ca. 15 seconds). The data were normalized using an empirical fit of each curve with a sum of two exponential functions and a linear drift, which describes the data very accurately (Figure S4A). The total amplitude changes with concentration, approaching zero (no LLPS) below 1 µM, in excellent agreement with the p53 concentration onset for LLPS observed by fluorescence microscopy (Figure S4B and Figure 1B). The kinetics of condensation may present a lag phase, depending on the conditions and the values of the rate constants (53). No lag phase was observed for p53 condensation under these conditions (Figure 3A). Normalization of the curves was calculated to better appreciate the changes in the half-time for completion of the process, *t*_1/2_, using an empirical fit of each curve with a sum of two exponential functions and a linear drift (Figure S4A). The *t*_1/2_ is strongly dependent on ionic strength, increasing 4-fold at sodium chloride concentrations between 20 and 40 mM (Figure S4C).

**Figure 3.**
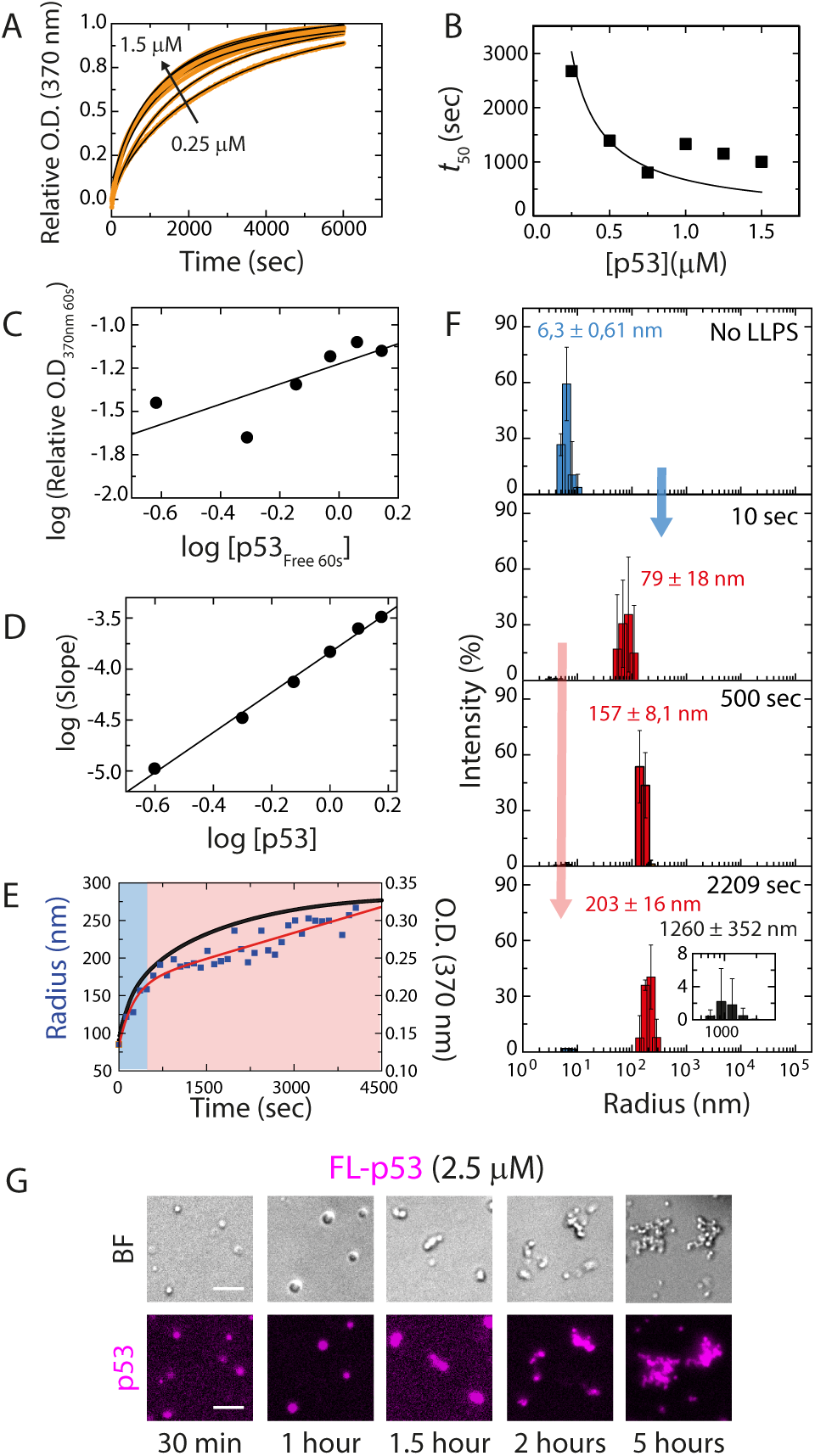
Kinetics of p53 condensation. (A) Time course of the kinetics at initial p53 tetramer concentrations from 0.25 to 1.5 µM. The data were normalized and fitted to the crystallization-like model from Martins and coworkers using the NAGPKin server (48) (See text). (B) The concentration dependence of the obtained *t*_1/2_ was used to extract the rate parameters *k*_α_ and *k*_β_. (C) Determination of the nucleus size following the model by Zlotnick (55). The nucleus size is calculated from the slope of the log-log plot of the concentration of dense phase versus dilute phase p53 tetramers after 60 seconds. According to this model, the observed nucleus size was 0.58 ± 0.24. (D) According to the Zlotnick model, the reaction order is calculated from the slope of the log-log plot of the initial rate (as in a linear fit to the first 110 seconds of the process) versus the initial concentration of p53 tetramer, yielding a value of 1.96 ± 0.07, indicating that addition of p53 molecules to the condensate is a second order reaction. (E) Comparison of size-progress measured as hydrodynamic radius using dynamic light scattering (Blue squares) and mass-progress measured by turbidity (Black trace). Single exponential fit is depicted in red. The blue background in the plot represents the nucleation-condensation reaction, whereas the red background depicts the early condensate growth. (F) Size-progress measured as hydrodynamic radius using dynamic light scattering. Top plot: FL-p53 in a non-LLPS condition (0.15 M NaCl). Other plots: FL-p53 under LLPS condition (40 mM NaCl) after 10, 500 and 2200 seconds. The arrows indicate the reactions described in (E), i.e., primary nucleation and growth (Blue) and early condensate growth (Red). (G) Representative fluorescence and bright field (BF) microscopy images of a sample containing 2.5 µM cy5-FL-p53 visualized at different time points, showing evolution of spherical homogenous droplets to larger coacervates. Scale bar on images = 10 µm.

We have aimed at rationalizing this data and gaining mechanistic insight using two complementary approaches to the early kinetics of p53 condensation. First, we made use of the model by Martins and coworkers for the crystallization-like assembly of a molecular condensate (53). This model was originally developed to describe amyloid fibril formation but may be used to study any association process of multiple molecules of the same chemical entity (48). The crystallization-like assembly model describes condensation by a general mechanism involving supersaturation-dependent nucleation and growth steps. The concentration dependence of condensation can be used to extract crucial information on primary and secondary nucleation (48), growth (48) and the presence of parallel pathways (48, 54, 55). The model considers the formation of a primary nucleus from molecules in the solution, growth of the primary nucleus and the formation of secondary nuclei on the surface of the primary nucleus. The analysis of experimental data quantifies the relative importance of the kinetic steps of primary nucleation, secondary nucleation and growth. We fitted the model as implemented in the NAGPKIN server (48) to normalized mass-progress curves (Figure 3A) and to the scaling of *t*_1/2_ versus the initial p53 tetramer concentration (Figure 3B). The model describes the data satisfactorily well, with a fitted value for the critical solubility (C_c_) of 0.19 µM. The fitted autocatalytic rate (*k*_a_) is 0.030 s^-1^, which corresponds to the sum of the rate constants for growth (*k*+) and secondary nucleation (*k*_2_). Mass-progress curves alone are unable to distinguish between these two processes, which requires a detailed analysis using multiple size-progress curves at different concentrations (48). The dimensionless nucleation rate (*k*_β_=*k*_n_/*k*_a_) is 2 x 10^-9^ (48). This value is smaller than 0.1, indicating that primary nucleation is slower than the sum of rate constants for growth and secondary nucleation (48). A global fit of the data yielded a r^2^ value lower than 0.95 (not shown), suggesting plausible parallel processes such as coalescence and/or off-pathway condensation events (48).

As a complementary approach, we made use of the model by Zlotnick and coworkers (49) for the kinetically limited assembly of a molecular condensate. This model was originally developed to describe the assembly of viral capsids but may be used to study any association process of multiple molecules. The kinetically limited assembly model describes condensation as a cascade of low-order association reactions, where a rate-limiting “nucleation” step is followed by faster elongation steps. The concentration dependence of condensation can be used to extract useful information on both the nucleation and the elongation steps. The size of the nucleus can be calculated from the slope of a double logarithmic plot of the concentration of condensated versus free p53 tetramers at a fixed time point early in the reaction, typically at a time when the kinetics is well described by a straight line. In our case, we calculated the concentration of condensated and free p53 tetramers after 60 seconds (See Experimental Procedures). Figure 3C shows that the slope of the plot is 0.58 ± 0.24, which corresponds to a nucleus formed by a single p53 tetramer. We interpret that a slow early step in p53 condensation likely involves a conformational rearrangement of the initial p53 tetramer. The reaction order of the faster elongation step can be determined from the slope of a double logarithmic plot of the initial rate in the reaction (Typically at a time when the kinetics are well described as a straight line) versus the initial concentration of p53 tetramer (49, 56). In our case, we calculated the initial rates by fitting a straight line to the first 110 seconds of the process. Figure 3D shows that the slope of the plot is 1.96 ± 0.07, which corresponds to a second order step. We interpret that in a faster early step in p53 condensation p53 tetramers are added to the growing condensate one at a time.

Next, in order to determine the physical species along the condensation pathway, we tackled dynamic light scattering analysis (Figure 3E and 3F). We first measured the hydrodynamic radius (*R_H_*) of the p53 tetramer at 1.25 µM concentration and in ionic strength conditions that preclude condensation, yielding a value of 6.3 ± 0.6 nm (Figure 3F, top panel). After triggering LLPS by diluting p53 into 40 mM NaCl, the *R_H_* of the major species at 20 seconds is 79 ± 18 nm (Figure 3F, second panel from the top). This phase corresponds to the nucleation-condensation reaction described by the crystallization assembly model (*k*_a_, Figure 3B; Figure 3E, blue time window; and Figure 3F, blue arrow). The *R_H_* of the major species increases to 157 ± 8 nm after 500 seconds and to 203 ± 16 nm after 2200 seconds (Figure 3F, red arrow, and Figure 3E, red time window). We assign this phase to early condensates growth, and single exponential analysis of the evolution of the *R_H_* with time, shows an apparent rate (*k*_app_) of 0.004 ± 0.0002 s-1 (*t*_1/2_ 350 sec). Noticeably, the histograms are monodisperse, indicative of discrete sizes, compatible with nucleation-growth of independent nuclei/particles. The evolution of the size was also followed in detail in 10 second intervals by DLS, with the data shown in Figure 3E, overlayed to a mass-progress curve measured under the same conditions. The radius of the major species continues to grow up to 4500 seconds, the limit of the time range selected. At 2200 seconds, we observe small amounts of a species with a radius larger than 1 µm. This is beyond the size range limit of the technique and was not further analyzed (Figure 3F, inset in the bottom panel). We speculate that this is related to expected droplet coalescence events, compatible with the polydisperse nature of the particles (Figure 3F, inset). In line with this, analysis of the decanted condensates by microscopy (Figure 3G) show incomplete coalescence events leading to the bead-like coacervates starting at 1.5 hours (Figure 3G and see Figure S4D for more detailed analysis).

### Human papillomavirus E2 protein rescues p53 from the aggregation route

We have previously shown that the interaction of FL-p53 with the DNA binding domain of the human papillomavirus (HPV) E2 protein (E2C) led to immediate formation of stoichiometrically tuned heterotypic FL-p53-E2C condensates (36). Given that p53 loss-of-function is linked to its amyloid aggregation pathway, a target for therapeutic rescuing intervention, we decided to test a possible effect of HPV E2 on the aggregation route *in vitro*. To this end, FL-p53 was incubated for two hours to allow for the formation of the bead-like coacervates (Figure 4A), followed by addition of E2C to the pre-incubated FL-p53 sample at a 4:1 E2C dimer : p53 tetramer ratio, based on the heterotypic LLPS stoichiometry determined previously (36). As time elapsed, the FL-p53 two-hour incubated coacervates were reshaped into highly regular spherical droplets which gradually increased in size due to complete coalescence events (Figure 4A, left panel). Control experiments without the addition of E2C showed the clustered bead-like coacervates expected from time evolution of FL-p53 homotypic condensation during longer incubation periods (Figure 4A, right panel). Additional control experiments combining unlabeled proteins, cy5-FL-p53 with unlabeled E2C, or unlabeled FL-p53 with FITC-E2C showed that the process is not affected by either of the fluorescent labels used (Figure S5A, B and C). Based on these findings, we further added HPV E2C to FITC-FL-p53 incubated for 24 hours, observing after 30 minutes full re-shaping of the homotypic FL-p53 coacervates into large heterotypic droplets (Figure 4B). FRAP assays performed on p53 rescued into these newly formed heterotypic droplets showed an amplitude recovery of 85%, with a *t*_1/2_ of 33 seconds (Figure 4B). This higher diffusion of p53 molecules indicates lower viscosity for heterotypic droplets than for homotypic FL-p53 coacervates incubated overnight without E2C (*t*_1/2_ of 83 seconds, Figure 1H).

**Figure 4.**
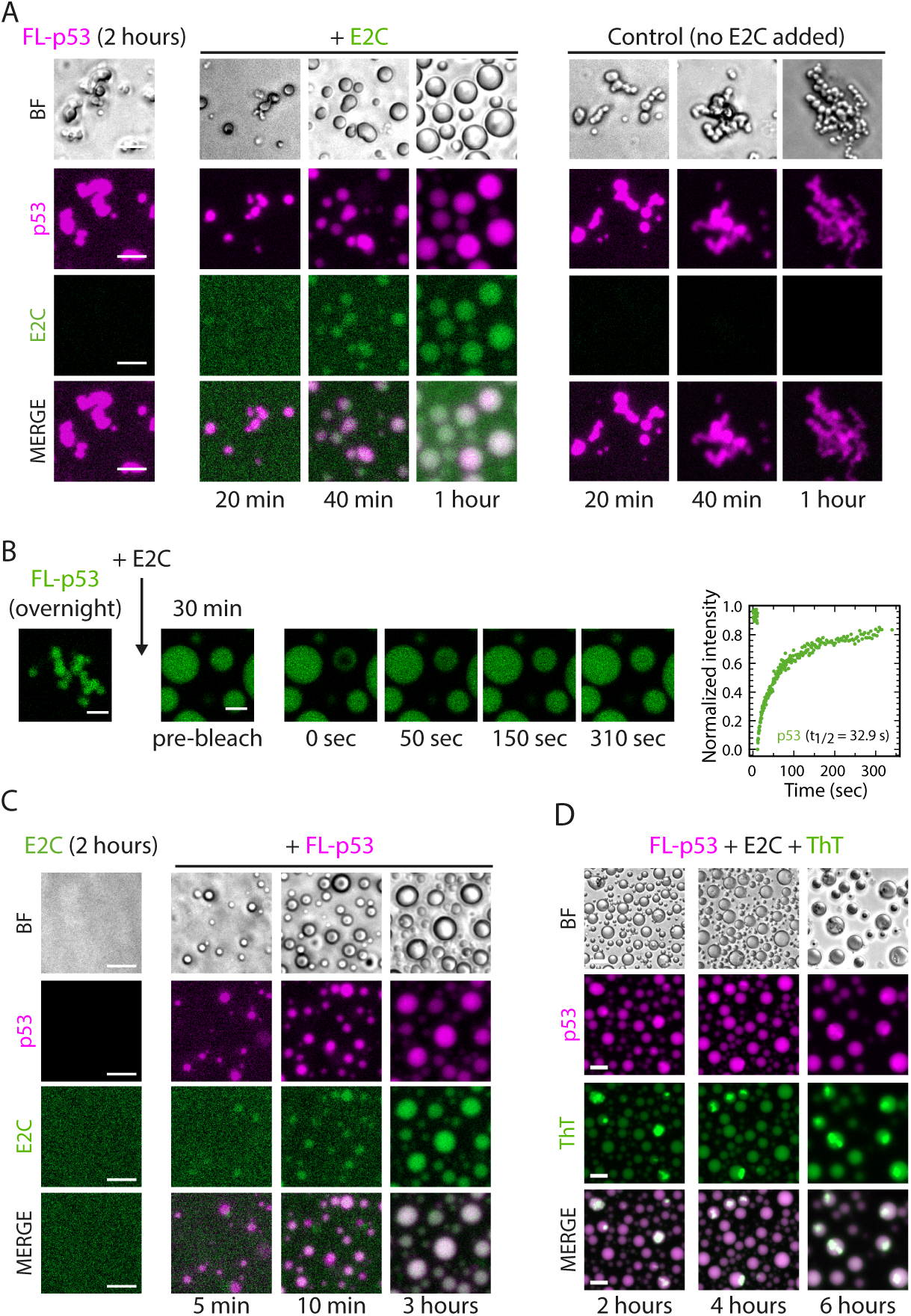
p53 aggregation fate is modulated by a viral protein. A) Representative microscopy images of a sample containing 2-hour incubated cy5-FL-p53 irregularly shaped coacervates (Left panel) followed by addition of FITC-HPV16 E2C (Central panel). Time-course examination of the evolution of the clusters into spherical droplets at 20 min, 40 min and 1 hour (Central panel). Control cy5-p53 homotypic samples without addition of E2C (Right panel). BF, bright field. Scale bars= 10 µm. (B) Representative confocal microscopy image of a FITC-FL-p53 sample incubated for 24 hours, showing irregularly shaped bead-like coacervates (Left image). Scale bar= 3 µm. Addition of HPV16 E2C (unlabeled) reshapes the clusters into large homogenous droplets, clearly visualized after 30 min post-addition of E2C to the sample. FRAP analysis was performed on these FITC-p53/E2C heterotypic condensates formed by reconstitution of the p53 homotypic coacervates (n= 2 droplets). Data were normalized to the average intensity of a droplet/region not photobleached and fitted using a double exponential function. Half time recovery (*t*_1/2_) of 32,9 ± 3.5 seconds was calculated. Scale bars= 3 μm. (C) Representative microscopy images of a sample containing 3-hour old FITC-E2C (Left panel) followed by addition of cy5-FL-p53 (Right panel). Time-course examination shows the assemble of a phase-separated system through the formation of heterotypic droplets. BF, bright field. Scale bars= 10 μm. (D) Bright field and fluorescence microscopy images of heterotypic droplets composed of cy5-FL-p53 and unlabeled E2C incubated with ThT during 2, 4 and 6 hours. BF, bright field. Scale bars= 20 μm.

In a different experiment, we reversed the order of addition of the proteins, incorporating FL-p53 without pre-incubation into a solution containing pre-incubated E2C. Homotypic FL-p53 droplets were formed within the first 5 minutes, the earliest time that allow for decantation required for microscopy (Figure 4C). This indicates that the immediate formation of these droplets was not affected by the presence of E2C in the solution (Figure 4C). Incorporation of E2C to the droplets starts to be detectable at 10 min and completed after three-hour incubation period (Figure 4C). We conclude that the formation of FL-p53 homotypic condensates is kinetically favored but shift to heterotypic condensates after the slow incorporation of E2C. The nature of the barrier for this event requires further investigation.

To gain insight into the amyloid-like behavior of p53 within the E2C rescued heterotypic droplets, we co-incubated a cy5-FL-p53/E2C mixture for 20 minutes with ThT and followed the time course by fluorescence microscopy (Figure 4D and Figure S5D). At two-hour incubation time, we observed diffuse homogeneous ThT staining in the droplets, matching the FL-p53 homogeneous distribution within the droplet (Figure 4D and Figure S5D). Interestingly, as time progressed a strong signal was detected in ThT-bound FL-p53 aggregates that partitioned heterogeneously within the droplets. These aggregates remained within the droplet boundaries, indicating that they are contained within the dense phase. The number of droplets containing these partitioned aggregates increased with time (Figure 4D), suggesting that despite a marked delay of the aggregation route by E2C, FL-p53 ends up accumulating as β-sheet-rich aggregates with ThT binding properties inside the droplets. Control experiments confirmed that the fraction aggregated is mainly composed of FL-p53, and not E2C (Figure S5E). Samples prepared under identical experimental conditions but in the absence of ThT allowed us to rule out any unspecific effect of ThT on the heterotypic LLPS system (Figure S5F).

### DNA dissolve p53 homotypic condensates

The fact that p53 is a transcriptional regulator and is present in the nucleus, prompted us to investigate the effect of DNA on the homotypic condensation process. First, we incubated FL-p53 for 20 minutes in the absence of DNA, observing the expected small regular condensates (Figure 5A, left panel). We added increasing concentrations of a 20 bp DNA duplex containing a high affinity consensus p53 binding site (46), that we name DNA_p53_. Sub-stoichiometric amounts of DNA_p53_ (0.25:1 DNA:p53) produced deformation of the droplets which were ultimately completely dissolved at exactly a 1:1 ratio (Figure 5A). These results indicate that the homotypic condensation of FL-p53 relies on a self-interaction that involves its DNA binding site, which is stoichiometrically saturated by DNA due to its high binding affinity (*K*_D_ in the range of 90 to 100 nM) (43). We next evaluated the effect of long stretches of DNA that better approximates the DNA encountered by the proteins within the nucleus. For this, we made use of calf thymus DNA (ctDNA), consisting of fragments of an average 1000 base pairs and variable sequence as it would be present in the nucleus. For the analysis of the stoichiometry, we consider 20 base pair non-specific sites along the entire ca. 1 kbp fragment, approximately 50 sites. Sub-stoichiometric amounts of ctDNA deformed the homotypic FL-p53 droplets and gradually led to their dissolution (Figure 5B). A residual of FL-p53 droplets observed may be due to the fact that the molar ratio of DNA is not high enough to completely dissolve FL-p53 phase separation but can also represent coexisting species of droplets that are more rigid and not affected by non-specific DNA, or that the inner dense phase is not accessible to a 1000 bp DNA.

**Figure 5.**
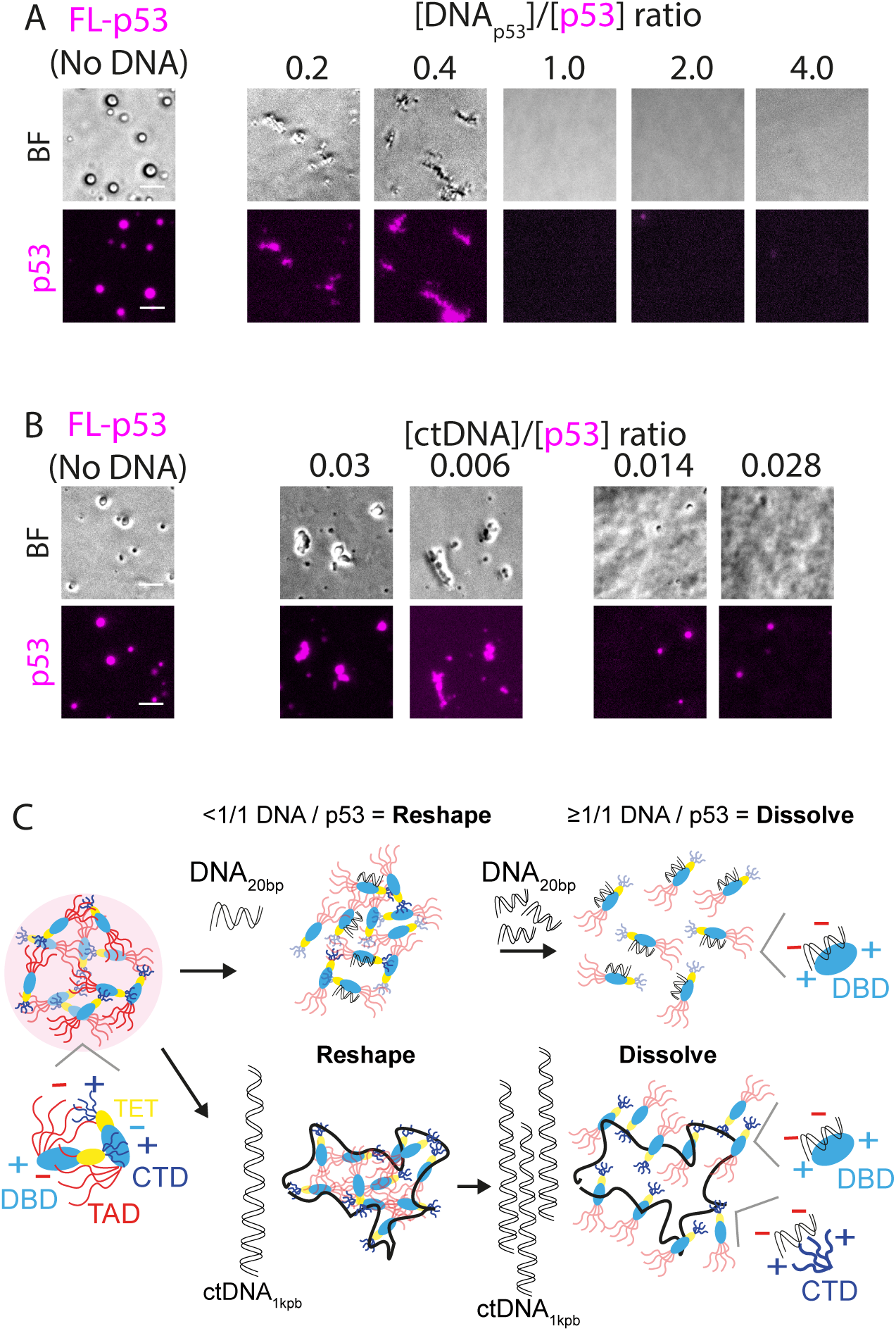
Effects of short specific and long unspecific DNAs on LLPS. (A) Representative microscopy images of samples containing increasing concentrations of specific 20 bp dsDNA for p53 (DNA_p53_) and 2.5 μM cy5-FL-p53. Control sample without DNA contained 2.5 μM cy5-FL-p53. BF, bright field. Scale bars = 10 µm. (B) Representative microscopy images of samples containing pre-formed p53 droplets followed by addition of increasing concentrations of calf thymus DNA (ctDNA). Control sample without DNA contains 2.5 μM cy5-p53. BF, bright field. Scale bars = 10 µm. (C) Model scheme of the effect of DNA on p53 LLPS. In the absence of DNA, the multiple interdomain interactions allows phase-separation of FL-p53, including its DNA recognition sequence in the DBD. When added, the p53 20 bp binding site duplex (DNA_p53-20bp_) diffuses into the droplets and reshape it to aggregates at sub-stoichiometric ratios. At over-stoichiometric ratios (>1:1 p53:DNA) the droplet is dissolved and all p53 is bound to DNA_p53-20bp_. ctDNA_1kb_ is also able to reshape the homotypic p53 droplets at sub-stoichiometric ratios, assuming ∼50 20pb long non-specific sites along the 1 kb fragment. In excess of sites, the droplets are dissolved.

## DISCUSSION

In this work we showed that a full-length pseudo wild-type variant of p53 can readily undergo homotypic LLPS, triggered rapidly by lowering the ionic strength. This results in small droplets of liquid nature in the absence of a crowding agent but large and regular droplets in the presence of PEG, and both processes are reversed by increasing the salt concentration. Deletion of the positively charged CTD abolished phase separation under the conditions that were permissive for FL-p53 and p53ΔTAD LLPS, in the absence of crowder, suggesting a main role for this domain. However, the p53ΔCTD can phase separate in the presence of PEG, suggesting that the TAD may also participate in LLPS even if its contribution is not as strong as the CTD. We have to consider that in the confined cellular environment, concentration of p53 in a condensate should be much higher, suggesting that both domains may contribute to homotypic condensation. In line with this, both TAD and CTD were shown to interact intramolecularly with the DBD (4, 6, 57), and the ampholytic nature of the protein and ionic strength sensitivity suggest that intermolecular interactions are likely to take place through the same domains that interact intramolecularly, providing a high valency that favors the network of weak interactions required for LLPS. In any case, both TAD and CTD partake in LLPS in the context of the intermolecular interaction network within the condensate. A study reported that neither TAD nor CTD deleted variants participate in LLPS, but we believe the discrepancy is likely caused by the use of a 17 mutation p53 protein in this study (32). Another study reported discrete LLPS for all FL-p53, p53ΔTAD, and p53ΔCTD, only in the presence of high crowder concentrations, in this case using an eGFP fusion not likely to be conformationally neutral for the native p53 ensemble in solution (39). We justify the use of the pseudowild-type mutant since the mutations are located in the core of the folded DBD and stabilize it. However, they do not affect the charge, and no mutations reside elsewhere, particularly in the intrinsically disordered domains (43).

The monomeric p53DBD protein was not able to phase separate under conditions that the FL-p53 does spontaneously, indicating not only the involvement of the substantial disordered regions (50% of the protein), but mainly that the tetrameric state of the protein is strictly required for droplet formation by providing the required multivalency (37, 58). The p53DBD monomeric protein was thoroughly studied on its ability to form fiber-like amyloid material by modifying the solvent conditions (59–61), and its participation in the amyloid aggregation process was confirmed by a short peptide within the DBD that disrupts aggregation and rescues the aggregated species (16, 62, 63). The isolated DBD was also shown to undergo LLPS in presence of high crowder conditions (15% PEG) (33). We conclude that the DBD could be determinant for the amyloid aggregation onset, but it cannot condensate on its own, something attainable only with the full-length protein that include the IDDs, the ampholytic sequences and, most importantly, tetramerization.

In this work, we have experimentally dissected the kinetic experimental condensation mechanism of p53, possibly one of the first protein LLPS systems to be tackled in an integrative manner. For a full description of the early p53 condensation reaction we can combine the results from mass-progress curves, size-progress curves, dynamic light scattering, and microscopy (Figure 6). p53 condensation takes place in a timescale of thousands of seconds and starts with formation of a nucleus from a protein tetramer. Subsequent steps involve a combination of formation of secondary nuclei and growth steps, where p53 tetramers are added to the growing condensate one at a time (Figure 6). Primary nucleation is slower than the sum of rate constants for growth (*k*_+_) and secondary nucleation (*k*_2_), which is consistent with the FRAP kinetics (akin to growth of a preformed condensate) being 10-fold faster that the condensation kinetics (Figures 1H and 3A). Kinetic modelling indicates that the condensation process is more complex than a slow nucleation followed by fast growth, with parallel processes such as coalescence and/or off-pathway condensation contributing significantly. An increase in ionic strength slows down the kinetics, suggesting formation of favorable electrostatic interactions in the transition states for the condensation reactions. The hydrodynamic radius of the main species becomes larger over time, as expected for growth of a condensate. Larger species appear in the dynamic light scattering and microscopy experiments on a slower timescale and are likely related to coalescence events. Further cutting-edge techniques are required to define the nature of the pre-coalescence condensate intermediates, including possible layered structural arrangements that transition to liquid as these coalesce into macroscopic droplets.

**Figure 6.**
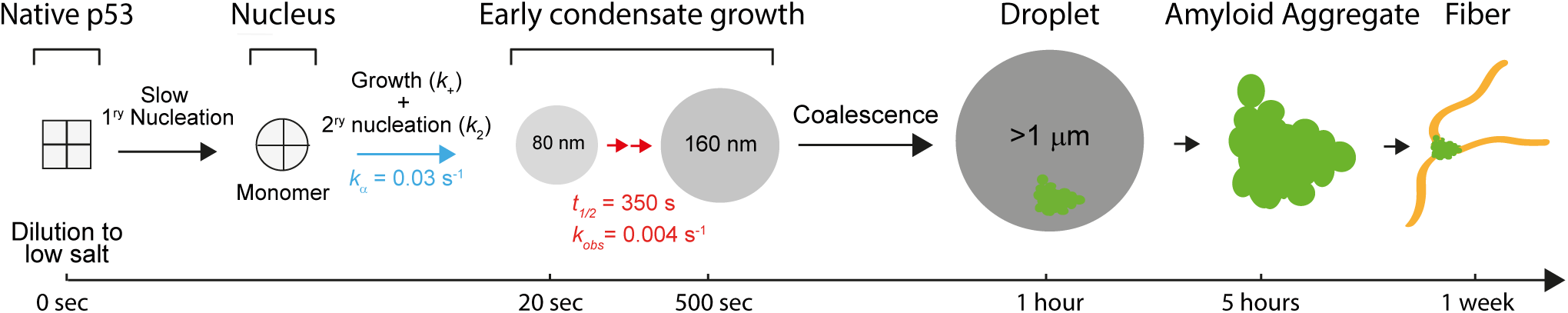
Integrated mechanistic model of p53 condensation-aggregation. Upon triggering condensation by dilution of the protein into low ionic strength buffer (see Figure 1), the native tetrameric ensemble of p53 undergoes a conformational change to yield the monomeric primary condensation nucleus, which further grows by gradual addition of p53 tetramers and secondary nucleation events (Blue, see Figure 3E and 3F) leading to early condensates. These condensates grow from 80 to ca. 200 nm *R_H_* monodisperse species (Red, Figure 3E and 3F) that coalesce into macroscopic droplets forming bead-like coacervates, with strong ThT binding (Figure 2A). These in turn slowly evolve into fibers with a branching growth pattern after a long incubation period (Figure 2E).

We showed that the homotypic p53 droplets slowly evolve to sticky bead-like coacervates, albeit of similar liquid properties than the initial droplets as judged by FRAP experiments (Figure 1H). Simple dilution experiments showed partial remodeling of these coacervates after 24 hours into droplets, strongly suggesting that the overall properties of these assemblies remain liquid-like (Figure 1G). More importantly, these coacervates show a strong binging of ThT as indicated by the fluorescence accumulation within the dense phase, starting at five hours and remaining in the one-week incubated species. Analysis of the week’s evolved coacervates clearly shows the presence of fiber-like materials, most noticeable by AFM. Fibrillar material of different lengths and regular 5-8 nm height emerge in a branching pattern from visible nuclei, suggesting a growth mechanism. These structures evolving sequentially from LLPS droplets to ThT binding coacervates to fibers, are not observed i) in conditions that abrogate LLPS (0.2M NaCl), ii) for the monomeric p53DBD construct, and iii) in the p53ΔTAD deletion. One study explored a possible LLPS-amyloid transition in full-length human p53 by the formation of stable transmissible cytoplasmic amyloid-like aggregate in yeast (41). Another study showed ThT-bound p53 aggregates in protein samples incubated at 37°C and proposed a non-classical fibril nucleation mechanism (34). These and our results do not rule out that the amyloid aggregation of p53 takes place though intermolecular interactions of the DBD region (33, 59), but clearly indicate that other regions are determinant for LLPS, and are required for both LLPS and aggregation routes of the full-length functional protein. Indeed, our fluorescence microscopy and TEM experiments indicate a functional segregation within the p53 molecule, with the C-terminal region being the LLPS driver, and the N-terminal region harboring key residues responsible for observed aggregation. Nevertheless, we do not know at this stage if there is a substantial degree of homogeneous structural order in the observed p53 fiber-like material.

In the last two decades, several therapeutic strategies directed to restore the native conformation of mutant p53 were addressed (18). The functional heterotypic condensation of FL-p53 with the C-terminal domain of the human papillomavirus E2 protein (E2C) (36) raised the question of whether the p53 conformations present in the matured homotypic coacervates could be rescued by E2C. We now showed that E2C progressively reshaped 2 hour- and 24 hour-aged coacervates into highly regular droplets, in what can be considered a time and E2C concentration dependent rescuing mode (Figure 4A, and B, Figure S5A, B, and C). While heterotypic p53:E2C droplets are formed in less than a minute after mixing (36), reshaping the 2-hour pre-incubated p53 homotypic coacervates by addition of E2C takes approximately 30 min to complete (Figure 4A, and B, Figure S5A, B, and C). The source of this energy barrier appears to be in the rate of incorporation of E2C to the dense phase of an otherwise matured homotypic p53 droplet (Figure 4A). Interestingly, the p53 rescued species in the heterotypic droplet exhibited a much more dynamic behavior than that of the coacervates (Figure 1H, and Figure 4B). This result confirms that “liquid-prone” conformations of p53 arise through interaction with E2C.

In the context of neurodegenerative disorders, it was reported that Hsp70 co-phase separates with FUS *in vitro*, co-localizes with FUS in stress granules (SGs), and, importantly, maintains the internal dynamics of FUS and other proteins to safeguard the liquid-like state of SGs (64). Also, nuclear import receptors (NIRs) were shown to inhibit cytoplasmic inclusions formed by aberrant phase separation transition and relocalize them to the nucleus in different RNA-binding proteins (RBPs), including FUS, hnRNPA1, hnRNPA2, and TDP-43 (65). In line with this, we previously showed that HPV E2C is able to relocalize mutant p53 to the nucleus (36). Therefore, as in the case of NIRs in neurodegenerative diseases (65), we propose that E2C might not only function as a modulator of LLPS, but also has the potential of a therapeutic agent in p53 cancer, starting from being a tool to investigate fundamental p53 rescuing mechanisms, that requires further exploration. Understanding whether small molecules designed for native restoration of stability p53 mutants are able to modulate the transition between p53 phase-separation and aggregation will be an important research area.

When assessing the amyloid-like behavior of the E2C rescued heterotypic droplets, although these remained highly regular and spherical after six hours of incubation, p53 only aggregates partitioned within the droplets. We propose that these are first condensed and distributed homogeneously within the droplet, and this in-droplet concentration increase subsequently evolve and partition to compact amyloid-like aggregates. The formation of sub-compartments within liquid BMCs has been reported (26, 66, 67). Since ThT binding structures partitioned within heterotypic droplets and did not evolve to large coacervates as observed for homotypic condensates (Figure 2A), we believe that within ageing heterotypic droplets, at least two compartmentalized populations exist. First, phase-separated complexes of p53 and E2C with homogeneous liquid behavior, the base for p53 rescuing activity of E2C. Second, p53 only amyloid-like aggregates, and we propose these are equivalent to the nuclei observed in AFM, from where the fibrillar material project and grow in a branching pattern (Figure 2E).

p53 is a transcription factor that operates within the nucleus where DNA is omnipresent. In this confined environment, tight-specific as well as non-specific DNA binding is expected. A minimal 20 bp DNA duplex with a specific p53 binding site completely dissolved the homotypic FL-p53 condensates at exactly 1:1 DNA_p53_:p53 ratio, clearly indicating that the high affinity stoichiometric binding and the excess of DNA shifts the equilibrium from LLPS to soluble p53-DNA complexes (Figure 5A). This result further supports the role of electrostatic interactions and particularly the crucial participation of the DNA binding site of the DBD (Model scheme in figure 5C), where DNA most likely competes with intra and intermolecule electrostatic interactions between the DBD and the acidic TAD (4, 68). A less plausible explanation of binding to the DBD-DNA recognition site as a main force for LLPS, is that p53 might undergo conformational changes induced by DNA that affect self-interaction through other regions.

Addition of the long ctDNA fragment disrupted the droplets but led to amorphous p53-DNA aggregates (Figure 5B). We speculate that multiple molecules of p53 bind multivalently and non-specifically either from the DBD or through the CTD, to the long DNA molecule, leading to aggregates instead of stable and fully soluble stoichiometric complexes as in the case of the short DNA duplex (Model scheme in figure 5C). In the presence of both high concentration of DNA (i.e., in the nucleus) and p53 (under stress), non-specific interactions will be the driving forces. Among the few studies assessing p53 LLPS in the cellular environment, most report p53 nuclear condensates that formed following DNA-damage (37, 39), whereas one study reported reversible cytoplasmic foci of p53 (41). It is therefore plausible, that homeostasis perturbation modulates the sub-localization of p53 condensates. Thus, in the cancer scenario, the cytoplasmic aggregates of p53 related to the dominant negative effect of p53 mutation (19), could arise from cytoplasmic condensation of non-transcriptionally active p53.

The strong tendency of p53 to undergo LLPS is based on several structural traits, i.e., modularity, tetrameric nature, nucleic acid binding, IDRs, charge distribution, ampholytic nature, and regulation by post-translational modifications (5, 69). We have shown that homotypic p53 LLPS is on-pathway to amyloid aggregation, that results at least in part in fibrillar material. This is a first report of otherwise elusive full-length p53 fibers (35) that require further investigation on their structure and homogeneity, not known at this stage. Further studies should address the nature of these fibers, together with the effect of hot-spot mutations, how this is translated to LOF and GOF effects, and how these impacts cellular proliferation in the various cancer types where p53 is mutated. Full overall understanding of the mechanism, species and key kinetic and thermodynamic steps along the pathway to pathological aggregation, including p53 conformers in solution, condensates, and amyloid aggregates, will uncover novel flanks for therapeutic conformational rescue in this validated drug target.

## AUTHOR CONTRIBUTIONS

S.S.B produced the plasmid constructs, produced and labeled all the recombinant proteins and prepared the DNAs, set up and performed the experimental work, analyzed the data, generated the figures, provided the funds for the project, and co-wrote the drafts and final form of the article. R.P.M performed the kinetic experiments and analyzed the data for the condensation mechanism. A.A.S helped with recombinant protein expression and purification. S.A.E performed and analyzed the TEM experiments and participated in generating the final form of the article. L.L performed and analyzed the AFM experiments. J.G.P performed and analyzed the TEM experiments. S.V, guided the TEM analyses and provided funds for the project. I.E.S designed and guided the mechanistic models and co-wrote the final form of the article. G.P.G conceptualized the project, guided the experimental work, provided the funds for the project, and co-wrote the drafts and final form of the article.

## ACKNOWLEDGMENTS

S.S.B, S.A.E, L.L, I.E.S, and G.P.G are CONICET career investigators. R.P.M is CONICET doctoral fellow. A.A.S is ANPCyT postdoctoral fellow. This work was funded by Agencia Nacional de Promoción de la Investigación, el Desarrollo Tecnológico y la Innovación (ANPCyT), grant PICT 2019-03295 for G.P.G. and grant PICT-2021-GRF-TI-0302 for S.S,B, and by Instituto Nacional del Cáncer (INC) grant INC S002460 for G.P.G.

