## Supplementary Figures 1-5 for "Experimental Kinetic Mechanism of P53 Condensation-Amyloid Aggregation"

Silvia S. Borkosky *et al.*

Figures S1 to S5

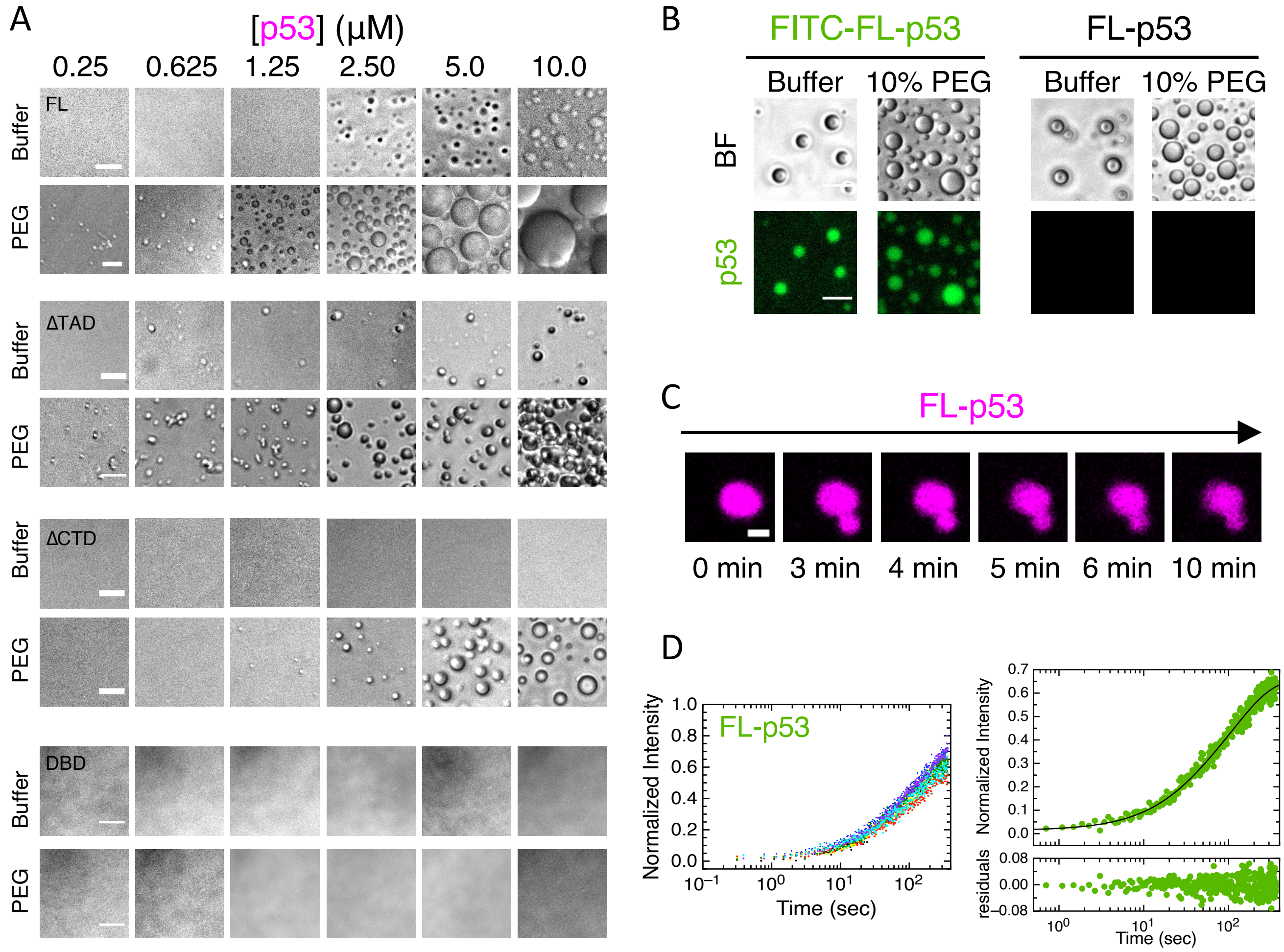


**Figure S1. Characterization of p53 homotypic LLPS.** (A) Representative bright field images of samples with increasing concentrations of cy5-labeled full-length p53 (FL-p53) and p53 truncated mutants, lacking the N-terminal TAD domain (ΔTAD), the C-terminal regulatory domain (ΔCTD) and the monomeric p53 DNA-binding domain (DBD), in buffer composed of 50 mM Tris-HCl, 40 mM NaCl, 1 mM DTT, pH 7.0. Spherical droplets are formed by FL-p53, which increase in size and number in accordance with protein concentration and are enhanced by molecular crowing. Samples with p53ΔTAD show presence of scarce spherical condensates in the absence of crowding, but irregular condensates in the presence of 10% PEG-4000. In the samples containing p53ΔCTD small discrete droplets were observed but only in presence of 10% PEG and at higher protein amounts. p53DBD remained soluble at all the concentrations evaluated and under both conditions. Scale bars= 10 µm. (B) Representative microscopy images of homotypic droplets formed in samples containing 2.5 µM FITC-FL-p53 (Left panel) or 2.5 µM unlabeled FL-p53 (Right panel) in absence or presence of PEG. BF, bright field. Scale bars= 10 µm. (C) Time-lapse confocal microscopy images of two homotypic cy5-FL-p53 droplets show incomplete coalescence leading to an irregularly shaped cluster. Scale bar= 2 µm. (D) Right, plot of FRAP analysis of FITC-FL-p53 homotypic droplets (n= 7 droplets) shows consistent replicates, indicating the reproducibility of the data.

Data were normalized to the average intensity of a droplet not photobleached. Left, normalized FRAP curve (Green dots) fitted with a double exponential function (Black line). The residuals are shown, justifying our choice of a double exponential function. A major recovery phase of 75% amplitude was obtained from the doble exponential fitting. A minor fast recovery phase was also observed but could not be explored further under the current experimental conditions.

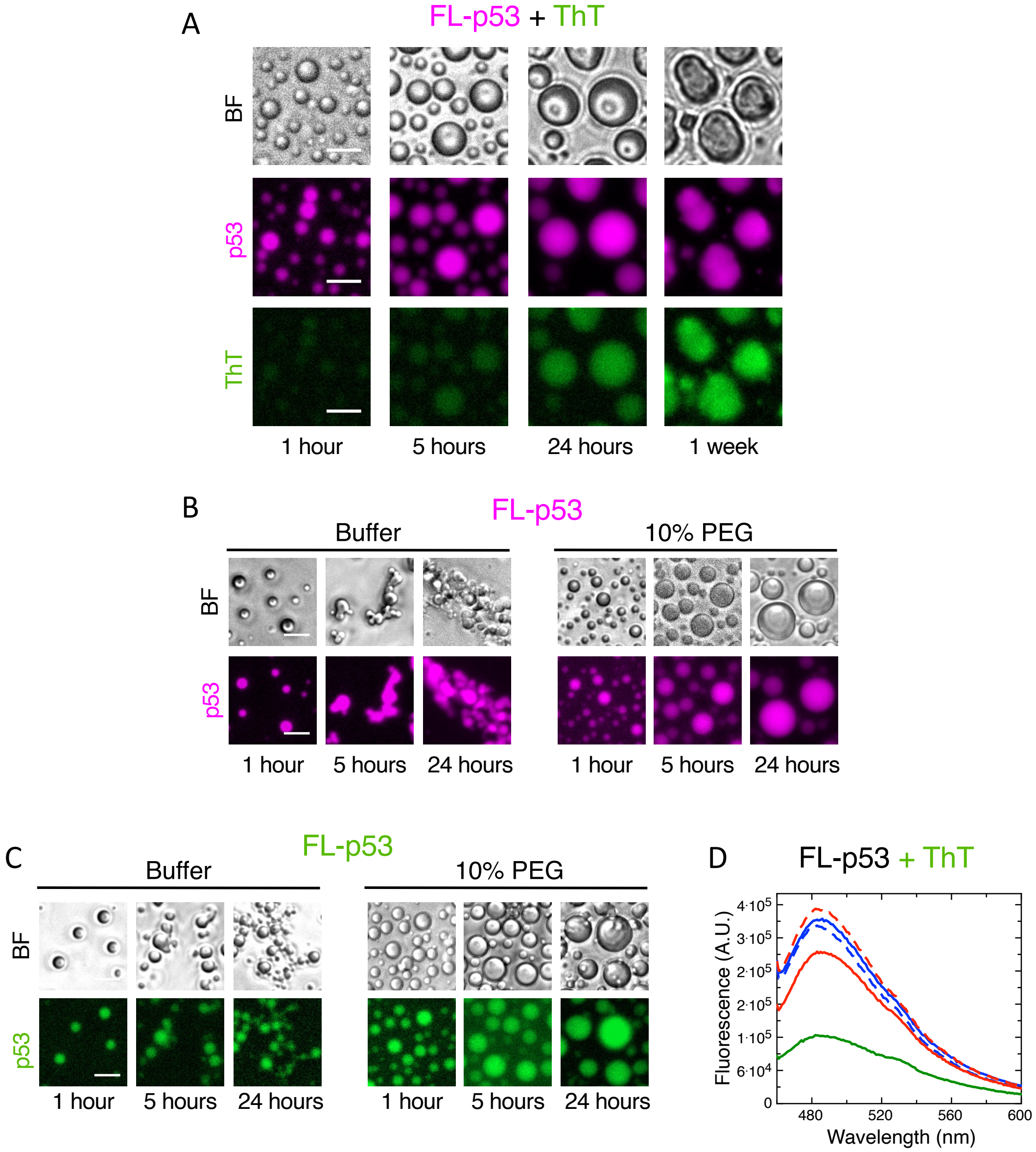


**Figure S2. Time-lapse evolution of p53 homotypic droplets.** (A) Representative microscopy images of 2.5 µM cy5-FL-p53 samples incubated with 50 µM Thioflavin T (ThT) during 1, 5, 24 hours, and 1 week in the presence of 10% PEG, used as crowding agent. BF, bright field. Scale bars= 10 µm. (B) 2.5 µM cy5-FL-p53 homotypic samples without addition of ThT (Control samples) showing time-course progression of spherical droplets into coacervates when crowder was not present (Left panel) or into larger regular droplets in presence of 10% PEG (Right panel). Scale bars= 10 µm. (C) p53 homotypic samples under similar conditions as in A but using FITC-labeled FL-p53. BF, bright field. Scale bars= 10 µm. (D) Fluorescence spectra of FL-p53 in buffer containing 25 µM ThT and 40 mM or 150 mM NaCl, pH 7.0, incubated for 1 hour (Thick line) or 1 week (Broken line). The spectrum depicting the fluorescence emission of buffer containing ThT without protein is shown in green.

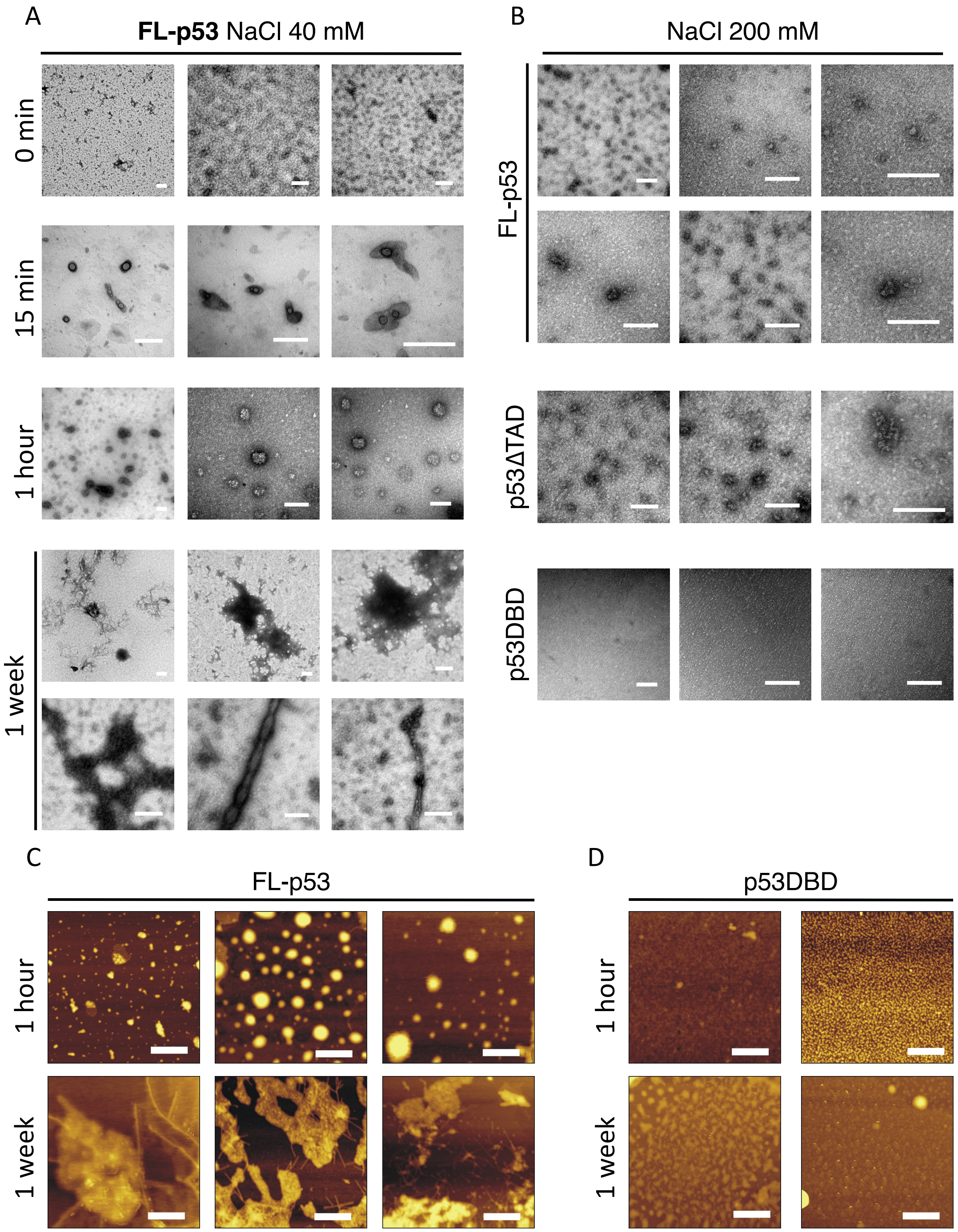


**Figure S3. Transmission electron microscopy and atomic force microscopy of p53 condensates.** (A) Samples containing 2.5 µM FL-p53 in 50 mM Tris-HCl, 40 mM NaCl, 1 mM DTT, pH 7.0 were visualized by transmission electron microscopy (TEM) at different incubation time-points. Species compatible with liquid-like droplets can be observed in samples incubated for 15 min and 1 hour. In FL-p53 samples incubated for 1 week, fibril-like and amorphous-like species can be appreciated. Bar on images= 200 nm. (B) Samples containing 2.5 µM FL-p53, p53ΔTAD, or p53DBD incubated for 1 week in buffer containing 50 mM Tris-HCl, 200 mM NaCl, 1 mM DTT, pH 7.0, show the absence of aggregated species in any of the p53 protein variants. Bar on images= 200 nm. (C) Atomic force microscopy (AFM) visualization was carried out to complement the results obtained by TEM. AFM images of FL-p53 samples incubated for 1 hour show the presence of spherical condensates in most of the observed fields, but also areas with irregularly shaped structures. One week incubation of FL-p53 under the same conditions show the presence of large irregular aggregated structures interspersed with fibrillar-like species. Bar on images= 500 nm. (D) AFM visualization of control samples of monomeric p53DBD in buffer composed of 50 mM Tris-HCl, 40 mM NaCl, 1 mM DTT, pH 7.0, show absence of aggregates, at both time-points evaluated. Bar on images= 500 nm.

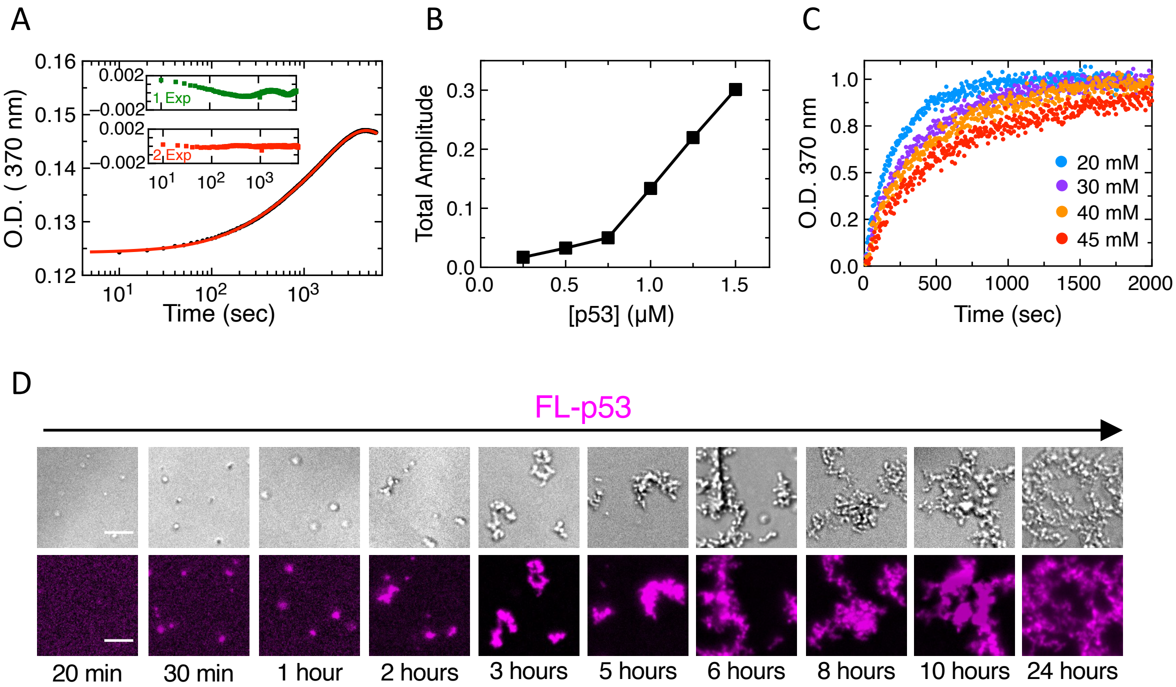


**Figure S4. kinetics of p53 LLPS.** (A) Kinetic turbidity traces of FL-p53 condensation were fitted to a sum of two exponential functions and a linear drift. Insets, residuals obtained with single or double exponential fits, justifying our choice of a double exponential fit. (B) Total amplitude of the turbidity signal at the different p53 concentrations were plotted as a function of initial tetramer concentration, showing clearly that amplitude approaches zero for concentrations below 1 µM. (C) Normalized p53 turbidity traces at NaCl concentrations ranging from 20-45 mM NaCl shows that $t_{1/2}$ of the condensation reaction increases with higher NaCl concentrations, suggesting a high dependance with ionic strength. (D) Representative microscopy images of a sample containing 2.5 µM cy5-FL-p53 visualized at different time points, showing evolution of spherical homogenous droplets to irregular condensates that fuse into larger coacervates. Note the size increase upon time elapse. BF, bright field. Scale bar on images= 10 µm.

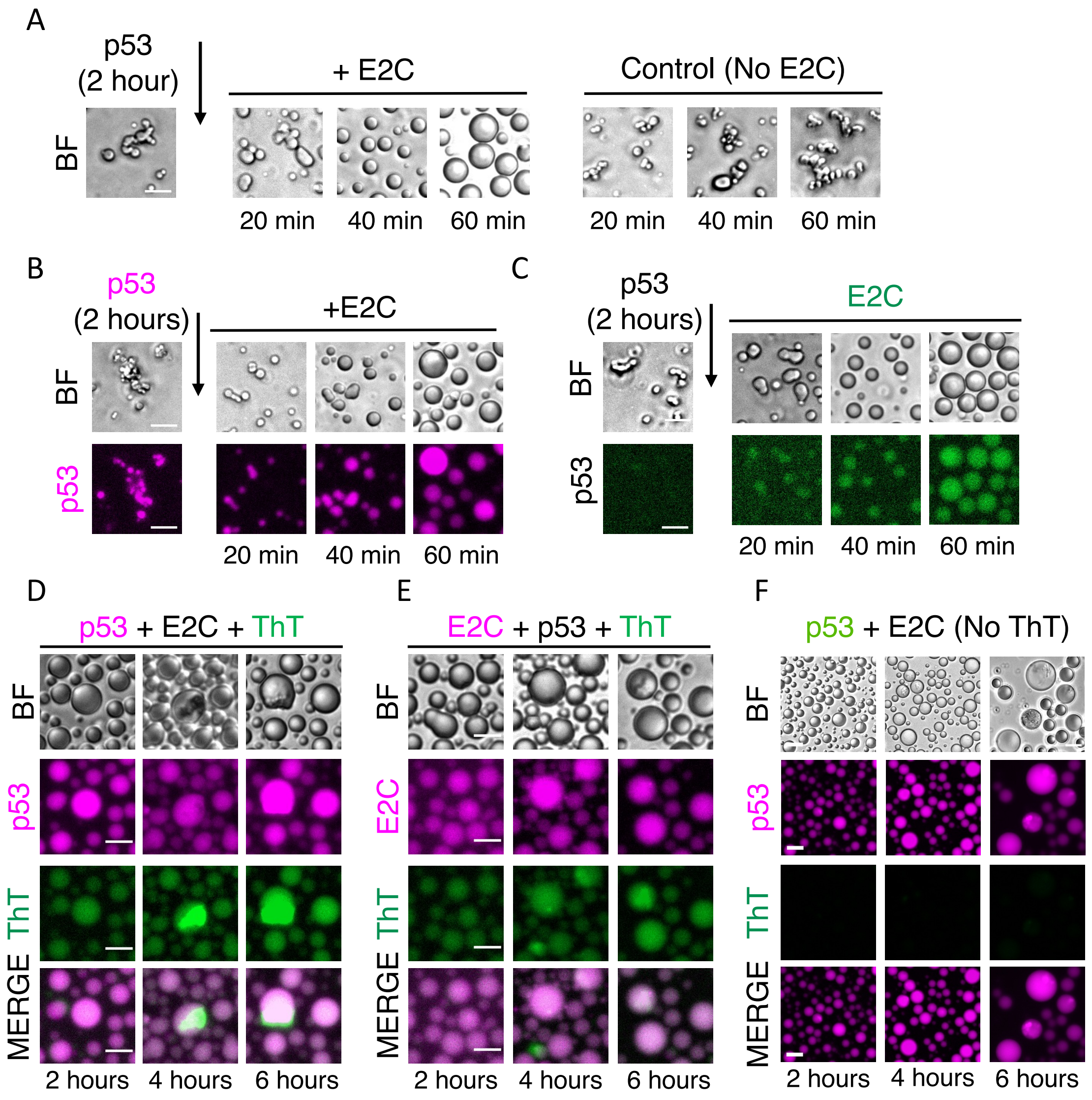


**Figure S5. E2C rescues p53 from the aggregation route.** (A) Representative bright field (BF) and fluorescence microscopy images of time-course examination of a sample containing 2-hour old FL-p53 irregularly shaped coacervates followed by addition of E2C, using unlabeled proteins. FL-p53 coacervates are gradually remodeled into spherical homogenous heterotypic droplets. Control samples, without addition of E2C, show evolution of the coacervates into larger clusters of irregular morphologies. Scale bar= 10 µm. (B) BF and fluorescence microscopy images of a sample containing 2-hour old cy5-FL-p53 irregularly shaped clusters followed by addition of unlabeled E2C. Scale bar= 10 µm. (C) BF and fluorescence microscopy images of a sample containing 2-hour old FL-p53 irregularly shaped clusters, followed by addition of FITC-E2C. Scale bars= 10 µm. (D) BF and fluorescence microscopy images of heterotypic droplets composed of cy5-FL-p53 and unlabeled E2C with ThT during 2, 4 and 6 hours, confirming the formation of aggregates composed of FL-p53 withing the heterotypic droplets. Scale bars= 10 µm. (E) A similar experiment as in D but using cy5-E2C and unlabeled FL-p53. BF, bright field. Scale bars= 10 µm. (F) Control samples of heterotypic droplets composed of cy5-FL-p53 and unlabeled E2C incubated without ThT (Control samples from figure 4D). Scale bars= 20 µm.
